# Capsaicin potently blocks *Salmonella typhimurium* invasion of Vero cells

**DOI:** 10.1101/2022.01.02.474733

**Authors:** Joseph A. Ayariga, Daniel A. Abugri, Balagopal Amrutha, Robert Villafane

## Abstract

As at 2021, the center for disease control (CDC) reported that *Salmonella* causes 1.2 million illness in the United States each year, with a mortality rate approaching 500 deaths per annum. Infants, the elderly, and persons with compromised immunity are the population with higher risk of mortality from this infection. At present there is no commercially available, safe and efficacious vaccine for the control and management of *Salmonella typhimurium* (*S. typhimurium)*. More so, *S. typhimurium* has been shown to develop resistance against most antibiotics used for treatment of the infection. Capsaicin, a bioactive compound from *Capsicum chinense (C. chinenses)* is undoubtedly one of the most widely used spice in the world. This heat producing compound is not only been used as food additive but have been demonstrated to possess unique properties that have pharmacological, physiological, and antimicrobial applications. In this work, the antimicrobial property of pure capsaicin or capsaicin extract against *S. typhimurium* is tested to determine the compounds effectiveness in *S. typhimurium* inhibition. Capsaicin extract showed potent inhibition of *S. typhimurium* growth at concentrations as low as 100 ng/ml, whereas pure capsaicin comparatively showed poorer inhibition of the bacteria. Furthermore, both capsaicin extract and pure capsaicin potently blocked *S. typhimurium* invasion of an animal cell line *in vitro*. Taken together, this work revealed that capsaicin might work synergistically with dihydrocapsaicin or the other capsaicinoids to inhibit *S. typhimurium* growth, whereas individually, capsaicin or dihydrocapsaicin could potently block the bacteria entry and invasion of Vero cells.

## 1.0 Introduction

Over the last 4 to 5 decades, the infections related to nontyphoidal *Salmonella* have increased and continue to be a significant global burden in health care systems in most countries **(1, 2, 3, 4, 5)**. These are generally foodborne. Most outbreaks of *S. typhimurium* are facilitated through the acquisition of new traits that enhance their adaptability and virulence **(6, 7)**. For instance, the emergence of multidrug-resistant (MDR) *S. typhimurium* DT104 has been demonstrated to be caused by the acquisition of the MDR gene via a plasmid-mediated process **(8)**. Specifically, the MDR-AmpC phenotype *S. typhimurium* and *S. newport* has been demonstrated to be exclusively plasmid mediated by plasmid transfer MDR genes **(9)**.

*S. typhimurium* is a gram-negative bacterium that causes gastroenteritis, bacteremia, and focal infections. The symptoms include high fever and diarrhea. It is currently treated with trimethoprim and sulfamethoxazole combination or with ciprofloxacin, azithromycin, or ceftriaxone **(10)**. However, the potential to develop resistance against these drugs is high. Most nontyphoidal *Salmonella* cause mild to severe, including life-threatening, infections **(11)**. In the early 2000, 600 deaths occur per year in the United States alone due to infections with nontyphoidal *Salmonella* serotypes **(12)**. The resistance of Salmonellae to antimicrobials has been demonstrated to correlate with increased risk of hospitalization, invasive illness, and death **(13, 14, 15, 16, 17, 18, 19)**.

Several published works are highlighting the importance of natural bioactive compounds against microbes **(20, 21, 22)**. Capsaicin, a naturally existing bioactive compound in *C. chinense* have received extensive scientific scrutiny, most works unraveled some of the mechanism of action of this compound and throws new light to its interesting and increasingly huge potential medicinal applications, with the majority of data indicating its potential antimicrobial potency **(23, 24, 25, 26, 27, 28)**. Capsaicin has been demonstrated to possess several pharmacological and physiological properties, for example, capsaicin has been shown to have anti-inflammatory and pain relieving **(29)**, anti-cancer and anti-tumor **(30, 31)** properties, also it has been proved to show positive cardiovascular and gastrointestinal effects **(32, 33)**. Dietary capsaicin has been demonstrated to protect cardiometabolic organs from dysfunction **(32)**. Studies underlying the pharmacodynamics of capsaicin revealed that peroxidase is directly linked to capsaicin metabolism, for which peroxidase oxidizes capsaicin **(34)**. Peroxisomes are cellular compartments where the turnover of some reactive anionic species and complex lipids take place. In the peroxisomal compartment, capsaicin could be degraded by the peroxisomal β-oxidation enzymes into acyl-CoAs which inturn serves as as substrate for the mitochondrial carnitine **(35)**.

In a study on the antimicrobial property of capsaicin, Manini et al., 2015, demonstrated that *Salmonella enteritis* (*S. enteritis*) did not develop any resistance against capsaicin treatment, and they showed that sublethal concentrations of capsaicin blocked *S. enteritis* from adhering to A549 monolayers cells, and significantly reduced cell-invasiveness **(28)**.

In another study, Omolo et al., 2018 showed that capsaicin from the fruits of *C. chinense* cultivars demonstrated bactericidal and antifungal effects **(27)**. They revealed that *L. monocytogenes* and *S. aureus* were more susceptible to capsaicin than *Salmonella* and *E. coli* O157:H7, and that *C. albicans* was the most susceptible to capsaicin **(27)**. Qiu et al., 2012 revealed that capsaicin provided protection for mice from methicillin-resistant *Staphylococcus aureus* infection **(36)**. Chatterjee et al., 2010, also demonstrated that capsaicin is a potent inhibitor of cholera toxin production in *Vibrio cholerae* **(37)**.

Capsaicin continues to attract major scientific interest specifically relating to its inhibitory potential against food-borne pathogens **(38)**, *Helicobacter pylori* **(39)**, *Pseudomonas aeruginosa*, and against MDR-ESBL producing Escherichia coli **(40)**.

The development of antimicrobial resistance is frequent with *S. typhimurium*. For instance, *S. typhimurium* DT104 has been shown to be resistant to cephalosporins, trimethoprim, ampicillin, chloramphenicol, quinolones, streptomycin, sulfonamides, and tetracycline **(41)**. Although the global survey of *salmonellosis* has been documented **(42)** relatively little is known about the interaction of this enteric bacterium and one of the bioactive compounds of a commonly consumed food spice called capsaicin. Thus, this work was framed to investigate the effect of capsaicin and capsaicin extract on the growth of *S. typhimurium in vitro*. Secondly, we investigated the ability of capsaicin or capsaicin extract in blocking the attachment and invasion of animal cells by *S. typhimurium*.

## 2.0 Materials and Methods

### 2.1 Reagents

All chemicals (Hexane, methanol –LC-MS (≥ 99.9%), water, and ethanol absolute proof (≥ 99.5%) were all high-performance liquid chromatography (HPLC) grade from Sigma Aldrich, MO, USA. Capsaicin and dihydrocapsaicin standards were purchased from Santa Cruz Biotechnology Inc., CA, USA. Dulbecco modified eagle medium (DMEM) and FBS were obtained from ATCC (Manassa, USA). The concentrations of the capsaicin and dihydrocapsaicin standards were evaluated using a stock solution of 6 mg/ml capsaicin and a stock solution of 5 mg/ml dihydrocapsaicin. The standards were dissolved completely 10:1 hexane-ethanol mixture and data was published in a previous communication **(31)**.

### 2.2 Animal cells and Bacterial Strain

Bacterial strains BV4012(LeuAam414 supE(gln). T. Poteete) is *S. typhimurium* strain without phage or extrachromosomal elements. It contains an amber mutation at the amino acid position 414, and this strain was obtained from Dr. Robert Villafane laboratory (Alabama State University, USA). The bacteria strain was grown and maintained in LB broth or LB agar, under 37 °C. Vero cells (ATCC, CCL 81) were obtained from BEI Resources, VA, USA. These cells were cultured and maintain in a T-25 cm^3^ flask using DMEM supplemented with 10% FBS and 1% penicillin-streptomycin-amphotericin, at 37°C, 5% CO_2_.

### 2.3 Description of Plant and Collection of Plant material

*Capsicum chinense* fruits were ground to a thick, semi-solid paste which had the characteristic red color as of the fruit. The resulted fine paste was mixed with hexane and eluted. The elute was air-dried to obtain powdered extract **(31)**.

### 2.4 Growth kinetics of S. typhimurium post capsaicin, or capsaicin extract administration

Growth kinetics of *S. typhimurium* were performed in 96 well-plates containing varying concentrations of pure capsaicin, capsaicin extract, or ampicillin. Concentrations used for pure capsaicin, capsaicin extract or ampicillin were 1 mg/ml, 100 µg/ml, 10 µg/ml, 1 µg/ml, 100 ng/ml and 10 ng/ml for pure capsaicin, capsaicin extract or ampicillin. Growth kinetics were carried out using a similar procedure previously published by Marini et al., 2015 with slight modifications. In summary, 300 µl of *S. typhimurium* (∼10 × 10^6^ CFU/mL) from each treatment were placed on microtiter plates, incubated for 24 h at 37 °C and read at OD600 at 1-h intervals using a SpectraMax® ABS Plus Microplate Reader. *S. typhimurium* grown in LB medium in the absence of capsaicin served as controls. All experiments were performed in triplicate.

### 2.5 S. typhimurium live/dead assay

To test the bactericidal property of capsaicin or capsaicin extract from *C. chinense*, we employed the live/dead assay by using SYBR Green I (Invitrogen, USA) and propidium iodide (Sigma–Aldrich, USA) following a protocol previously published by Magi et al., 2015, that made use of the two nucleic acid dyes that differ in their ability to penetrate bacterial cells. In short, *S. typhimurium* was grown overnight, and the overnight culture diluted to an OD600 of 0.2 using LB broth supplemented with varying concentrations of capsaicin, capsaicin extract, or ampicillin. An untreated sample served as a control. Treated samples were incubated at 37 °C, 5% CO_2_ for 60 min. Afterwards, samples were stained with 1 × SYBR Green I or with SYTO-9 and 40 µg/ml propidium iodide, at room temperature for 25 min covered with aluminum foil. Subsequently, cells were harvested on filters (Ø = 0.2 µm, Millipore, USA), and examined under an EVOS FLC microscope (Life Technologies).

### 2.6 S. typhimurium membrane integrity test

To evaluate the mechanism of action of capsaicin or capsaicin extract on *S. typhimurium*, we tested if the compounds acted by disrupting bacterial cell membrane that might lead to cell membrane lysis. To test this hypothesis, 100 µl of *S. typhimurium* cells grown to OD of 0.5 were pelleted by centrifugation and resuspended in 1X PBS, then followed with incubation with lysozyme, capsaicin, capsaicin extract, or ampicillin at concentrations 100 µg/ml or 10 µg/ml for 20 or 60 min. After the set time points of incubations, 50 µl of each treatment were mixed with Agarose gel loading dye (Trackit, Thermo Fisher Scientific, USA). Samples were run in 0.8% agarose gel and gel stained with ethidium monoazide bromide for 30 min. Following rinsing, the agarose gel’s bands were captured using ChemiDoc installed with Quantity One software.

### 2.7 Vero cell viability determination under capsaicin or capsaicin extract treatment

To investigate the effect of capsaicin on Vero cells, the cells were cultured in 96-tissue well plates containing DMEM media supplemented with 10% FBS. Capsaicin at 100 µg/ml, 10 µg/ml, 1 µg/ml and 0.1 µg/ml were mixed with the culture media and incubated for 8 hours. Control cultures did not receive capsaicin treatment. At the 8-h time point, AlamarBlue at 10% were added to the growing culture and allowed to incubate for an additional 4 h. Absorbance at 570 nm was read using Cytation 3 imaging/plate reader (BioTek, USA) and data plotted on a bar chart to evaluate the percentage cell viability.

### 2.8 Determination of Vero cell inhibition under capsaicin or capsaicin extract treatment

To determine the ability of capsaicin or capsaicin extract to inhibit the growth of Vero cells *in vitro*, the cells were cultured as described above, and following viability assessment using AlamarBlue, the percent cell inhibition was calculated as:

((the absorbance value of the control group) − (absorbance of the treated group))/(sum of both the control and the treated groups) × 100.

### 2.9 Anti-adhesion assay

Vero Cells lines at a density of 1 × 10^5^ were cultured in DMEM media supplemented with 10% FBS (all from Gibco, USA) in a 96-tissue culture plate (Corning Costar, Milano, Italy) at 37 °C in an atmosphere containing 5% CO_2_ for 24 hours for cells to attach. After 24 h, the media was removed, and 300 µl of *S. typhimurium* cells at an OD of 0.3 were added to the growing Vero cells, this was immediately followed with capsaicin, capsaicin extract or media control supplemented DMEM media only. Infection was allowed for 30 min following a similar procedure by Manini et al., 2015 **(28)**. Final concentrations of capsaicin or capsaicin extract in the culture media were 100 µg/ml and 10 µg/ml. Prior to infection of the Vero cells, the *S. typhimurium* was freshly grown to an OD of 0.5, harvested, and pelleted via centrifugation. Bacteria pellets were washed thrice with 1X PBS and resuspended in DMEM media to an OD of 0.3.

To investigate the ability of *S. typhimurium* to attach to the monolayer Vero cells, the wells containing the infected Vero cells, or the controls were washed 3 times with 1X PBS and lysed with chilled distilled water. 300 µl of the lysed cells were spread to LB agar plates and incubated overnight at 37 °C. The colony-forming units (CFU) were counted to evaluate the total adherent bacteria in each 300 µl of the lysate.

### 2.10 Anti-invasion assay

The anti-invasion capacity of capsaicin or capsaicin extract was evaluated by counting the viable bacteria that survived the antibiotic insults due to their intracellular presence in the host cell’s cytoplasm. To investigate the intracellular presence of *S. typhimurium* in infected Vero cells, infected monolayers were washed 3 times with 1X PBS as previously demonstrated by Manini et al., 2015 **(28)**. This was followed with the administration of penicillin (1 mL of bactericidal concentrations of penicillin (5 µg/ml)) and gentamicin (100 µg/ml), and the samples were allowed to sit for 2 h at 37 °C in 5% CO_2_. The treated monolayers were washed and lysed with chilled distilled water, then viable intracellular bacteria were assayed by plating 300 µl of the lysate on an LB agar plate and cultured overnight. Each assay was repeated thrice.

### 2.11 Statistical analyses

For all statistical data, values were derived from multiple measurements (from replicates of 3 experiments) and averaged. Differences between groups were assessed with paired Student’s t-test using OriginPro software. Values are reported as mean ±SEM. P-values ≤ 0.05 were considered statistically significant.

## 3.0 Results and Discussion

### 3.1 Growth kinetics of S. typhimurium post capsaicin, or capsaicin extract administration

In this study, administration of pure capsaicin or capsaicin extract isolated from *C. chinense* fruit demonstrated that both exert bacteria inhibitory activity against *S. typhimurium* with differing efficacy. Similar reports of capsaicin and dihydrocapsaicin showing antibacterial poperties have been published **(28, 43)**. The capsaicin extract however exerted relatively higher inhibitory activity against *S. typhimurium* than the pure capsaicin at similar concentrations (See **Figures 1, 2**, and **5)**. The lowest concentration of capsaicin extract that potently reduced *S. typhimurium* growth was 10 ng/ml whereas pure capsaicin could not effectively reduced the growth of the bacteria at concentrations of 10 µg/ml or lower **(Figure 2)**. In a control experiment using ampicillin at varying concentrations, a lower concentration of 10 ng/ml potently reduced the growth of *S. tyhimurium* **(Figure 3)**. As shown in **Figure 4**, staining of *S. typhimurium* with SYTO-9 and Propidium iodide (PI) indicated that *S. typhimurium* growing on culture media pretreated with 1 mg/ml of capsaicin extract showed higher *S. typhimurium* killing (indicated by the higher red fluorescence of the PI). The control group that received no treatment showed no red fluorescence **(Figure 4)** since the undamaged bacterial membrane showed green fluorescence, but those with damaged membranes shows red fluorescence. For immunofluorescent images of other concentrations of capsaicin, capsaicin extract treatment of the bacteria, see Supplementary **Figures 1, 2**, and **3**.

**Figure 1.**
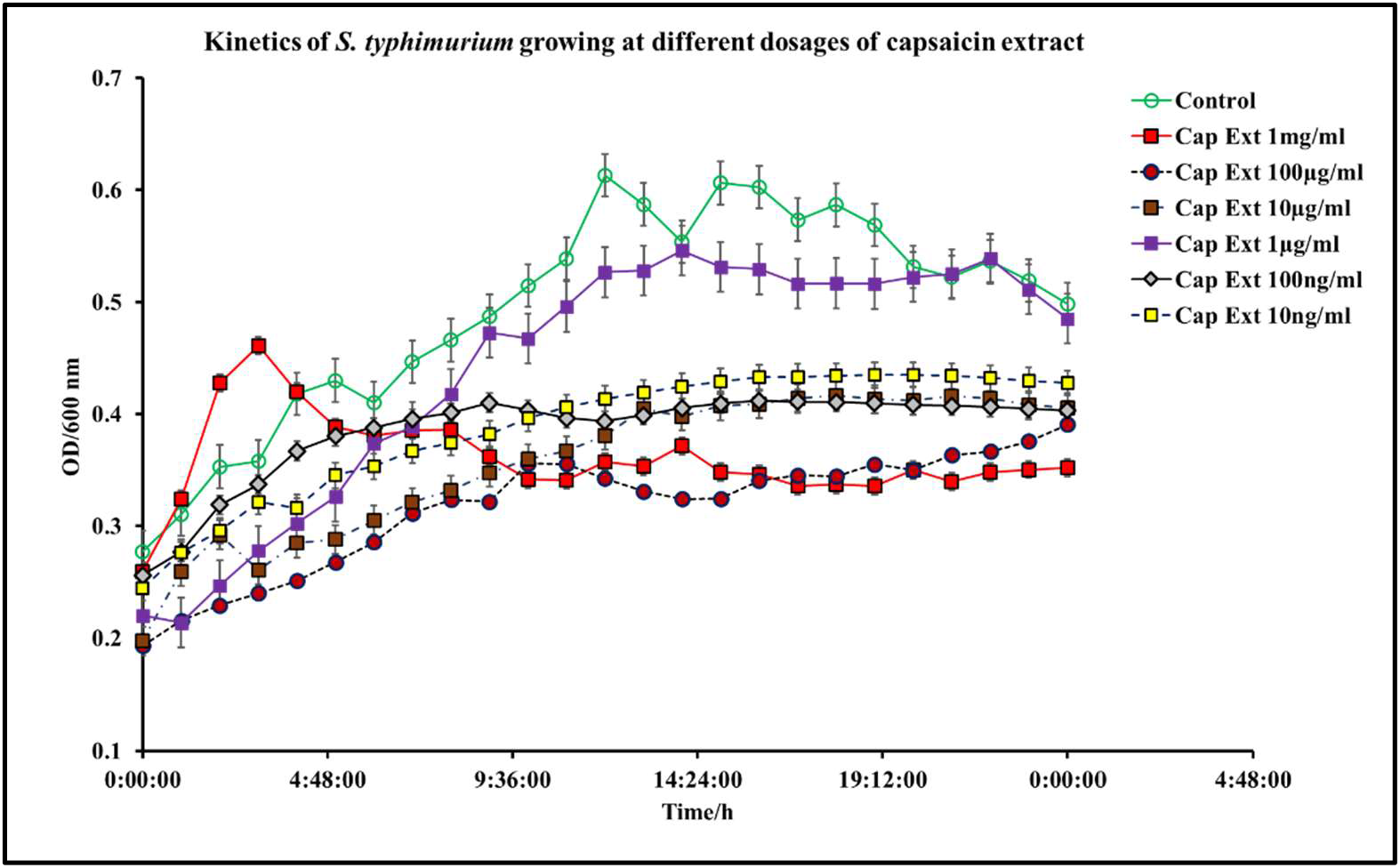
Kinetics of *S. typhimurium* growth at varying dosages of capsaicin extract. The results represent the average of three independent experiments ± standard deviation.

**Figure 2.**
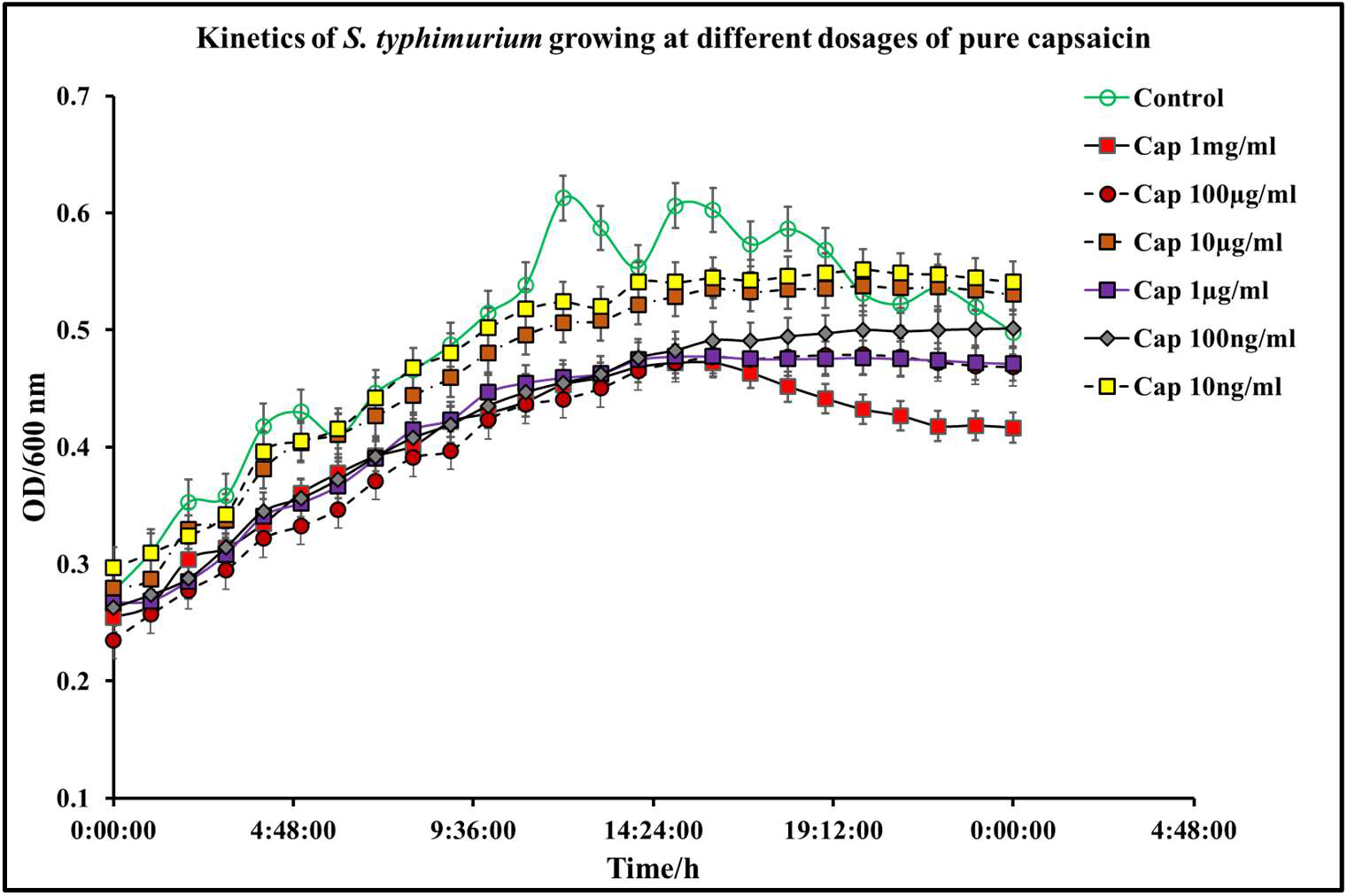
Kinetics of *S. typhimurium* growth at varying dosages of pure capsaicin. The results represent the average of three independent experiments ± standard deviation.

**Figure 3.**
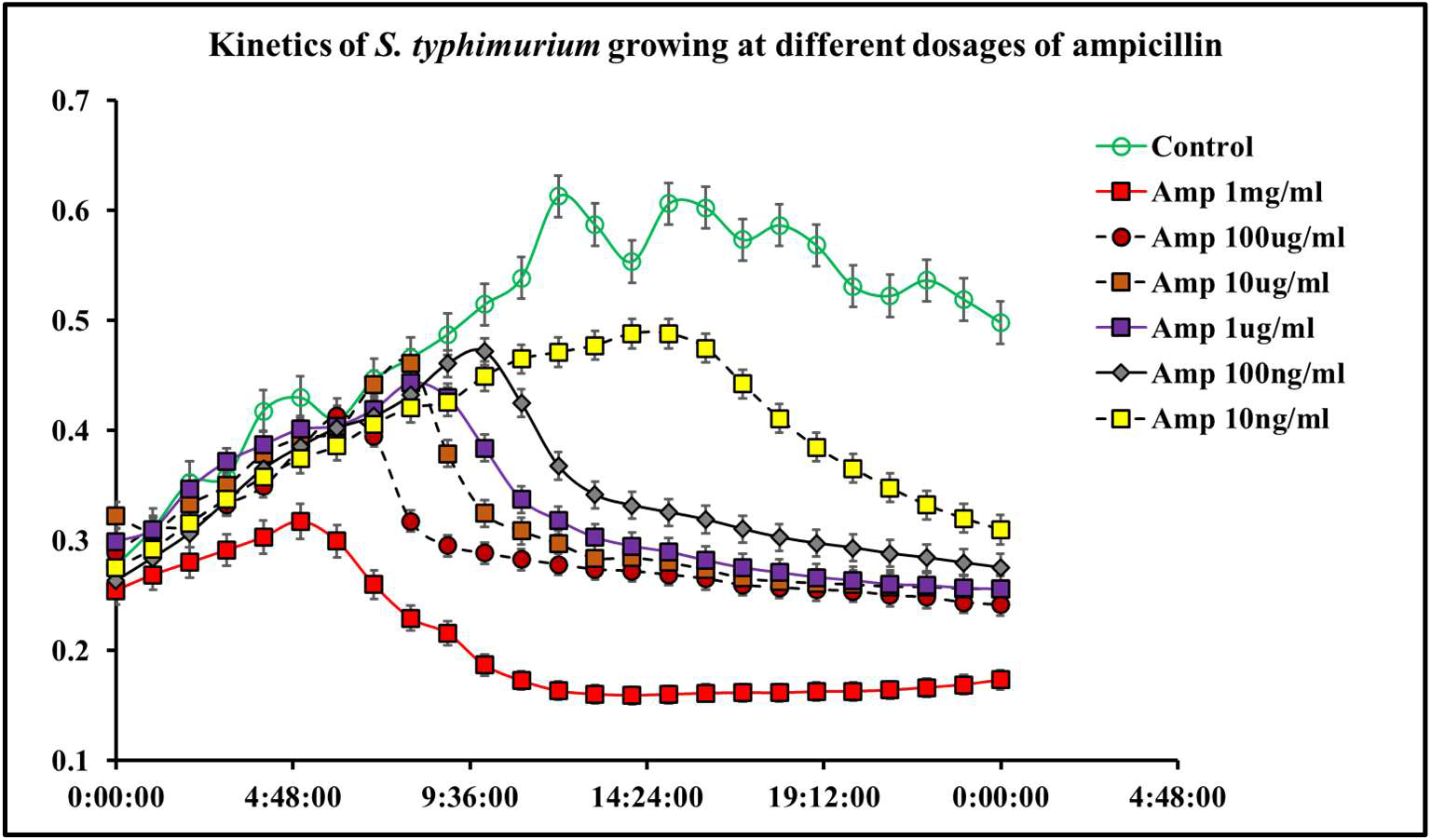
Kinetics of *S. typhimurium* growth at varying dosages of ampicillin. The results represent the average of three independent experiments ± standard deviation.

**Figure 4.**
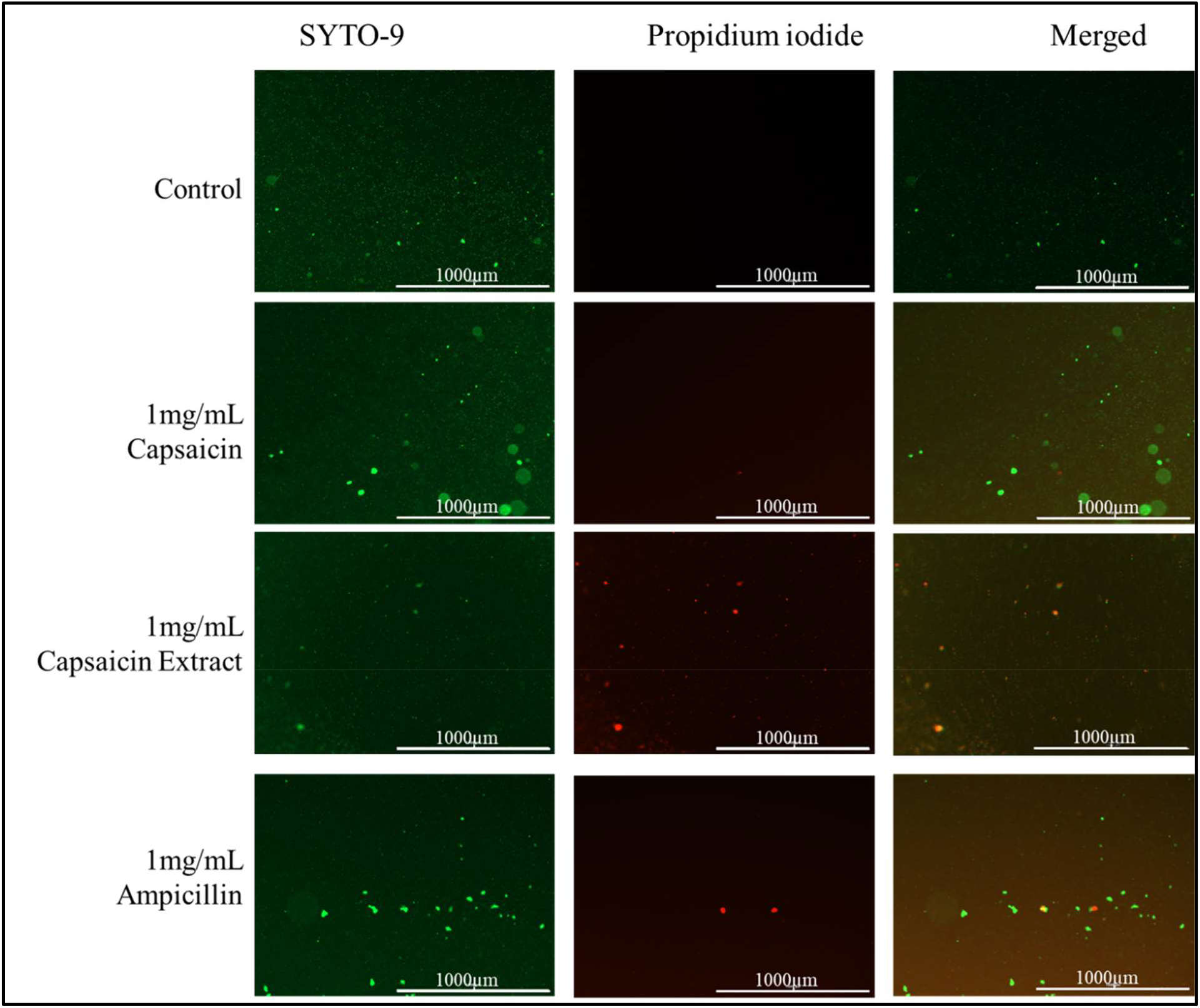
Immunofluorescent images of *S. typhimurium* growing on culture media pretreated with 1 mg/ml of pure capsaicin, capsaicin extract or ampicillin (30 min incubation time). Control received no treatment. Undamaged bacterial membrane shows green fluorescence, but those with damaged membranes shows red fluorescence.

**Figure 5.**
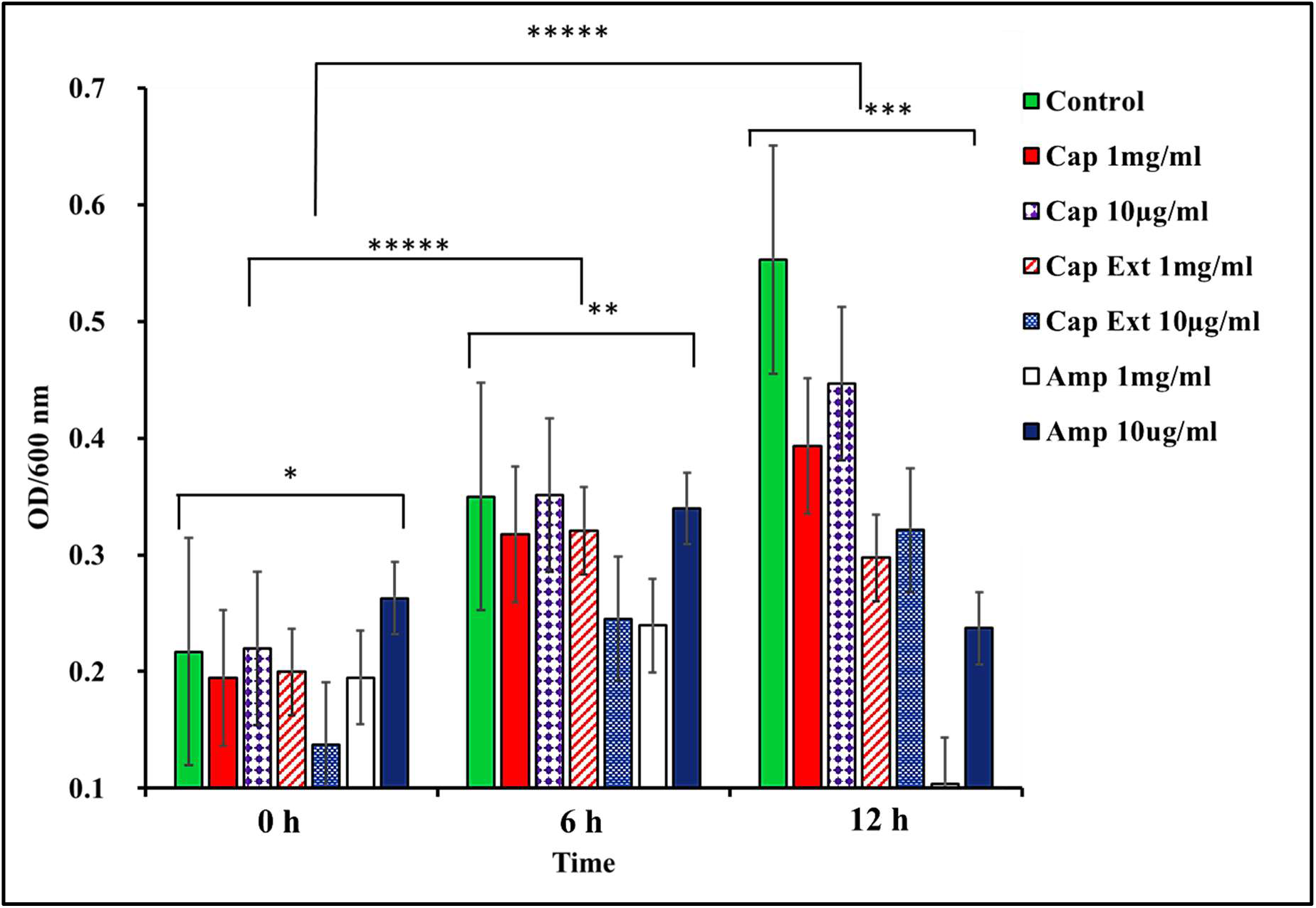
Comparing pure capsaicin, capsaicin extract and ampicillin ability to limit *S. typhimurium* growth at various culturing time points. * and ** p-value ≤ 0.1, ***, ****, and ***** p-value ≤ 0.05; n = 3.

As shown in Table 1, the IC_50_s of pure capsaicin and capsaicin extract were calculated for both 12 h and 24 h time points. Whereas 12 h period of pure capsaicin produced an IC_50_ of 34.0 µg/ml, this increased to 42.2 µg/ml at the 24 h time point. The capsaicin extract showed a lower IC_50_ of 23.1 µg/ml, but this value was increased to 90.2 μg/ml at the 24 h time point. These increases in IC_50_ values might be indicative of resistance development of the bacteria against both pure capsaicin and the capsaicin extract.

**Table 1.**
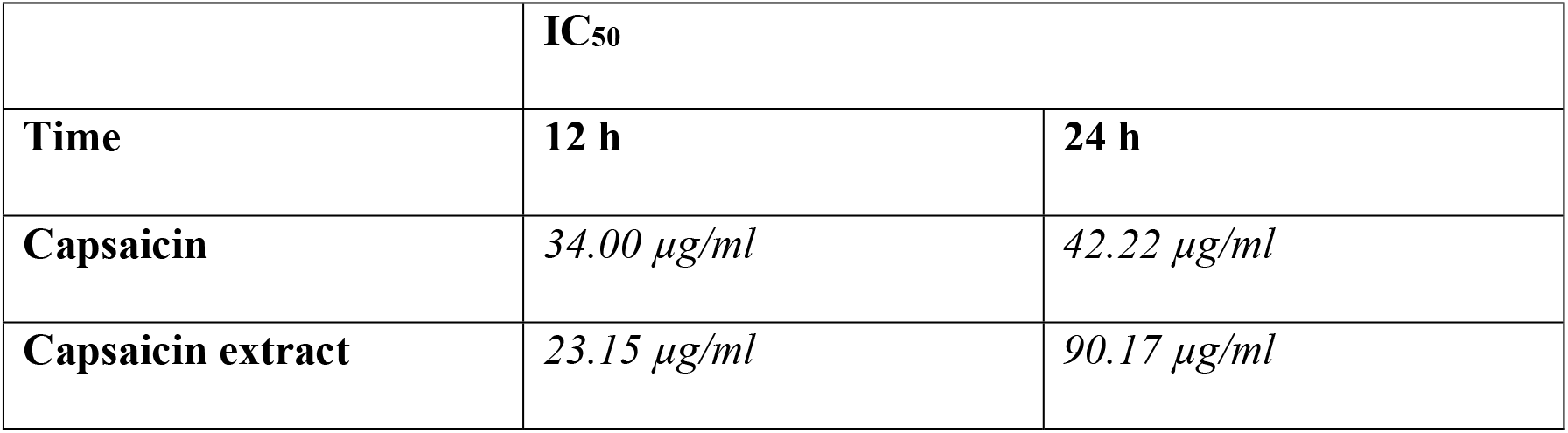
Median concentrations of bacterial growth inhibition (IC_50_s) produced by treatment with capsaicin or capsaicin extract. IC_50_ values are expressed in µg/ml with a 95% confidence. (n = 3; p-value ≤ 0.05).

### 3.2 Capsaicin promotes Vero cells growth at lower concentrations and poorly inhibits Vero cell growth at high concentrations

In this work, Vero cells were cultured in DMEM medium supplemented with varying concentrations of pure capsaicin or capsaicin extract. Viabilities of these cells were shown not to be significantly different (p ≤ 0.05). However, there were observed slightly higher growth of Vero cells at 0.1µg/ml of capsaicin than all other treatment groups **(Figure 6)**. Morphologically, no distinct differences were observed between cells of the different treatments groups of capsaicin, or capsaicin extract **(Figure 8)**. Infecting Vero cells with *S. typhimurium* showed that the bacteria attached to host Vero cells within 30 min post infection **(Figure 9)**.

**Figure 6.**
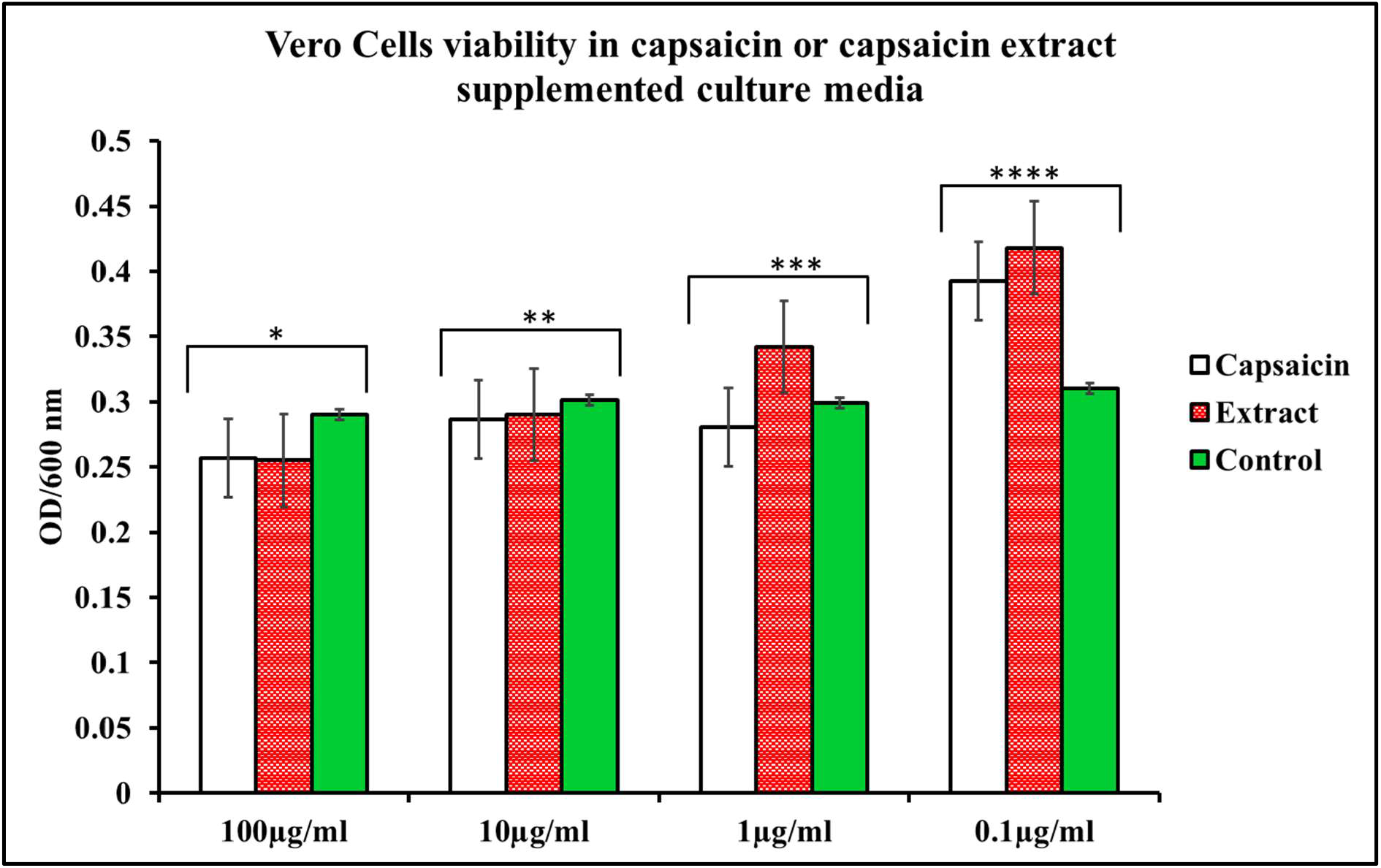
Viabilities of Vero cells in pure capsaicin or capsaicin extract at varying doses. *, **, *** and **** p-value ≤ 0.1; n = 3.

**Figure 7.**
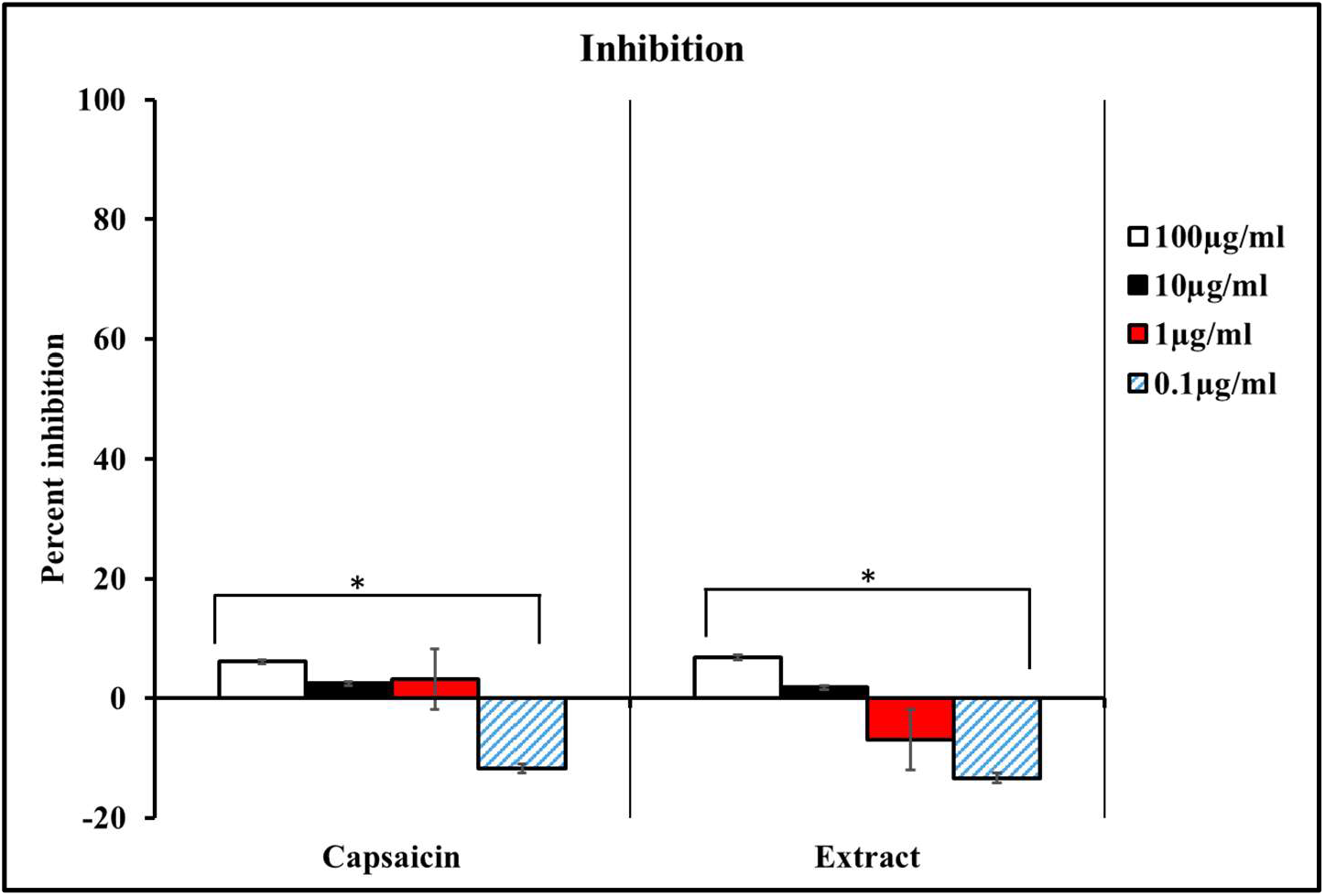
Percentage growth inhibition of Vero cells in pure capsaicin or capsaicin extract at varying doses. * p-value ≤ 0.05, n = 3.

**Figure 8.**
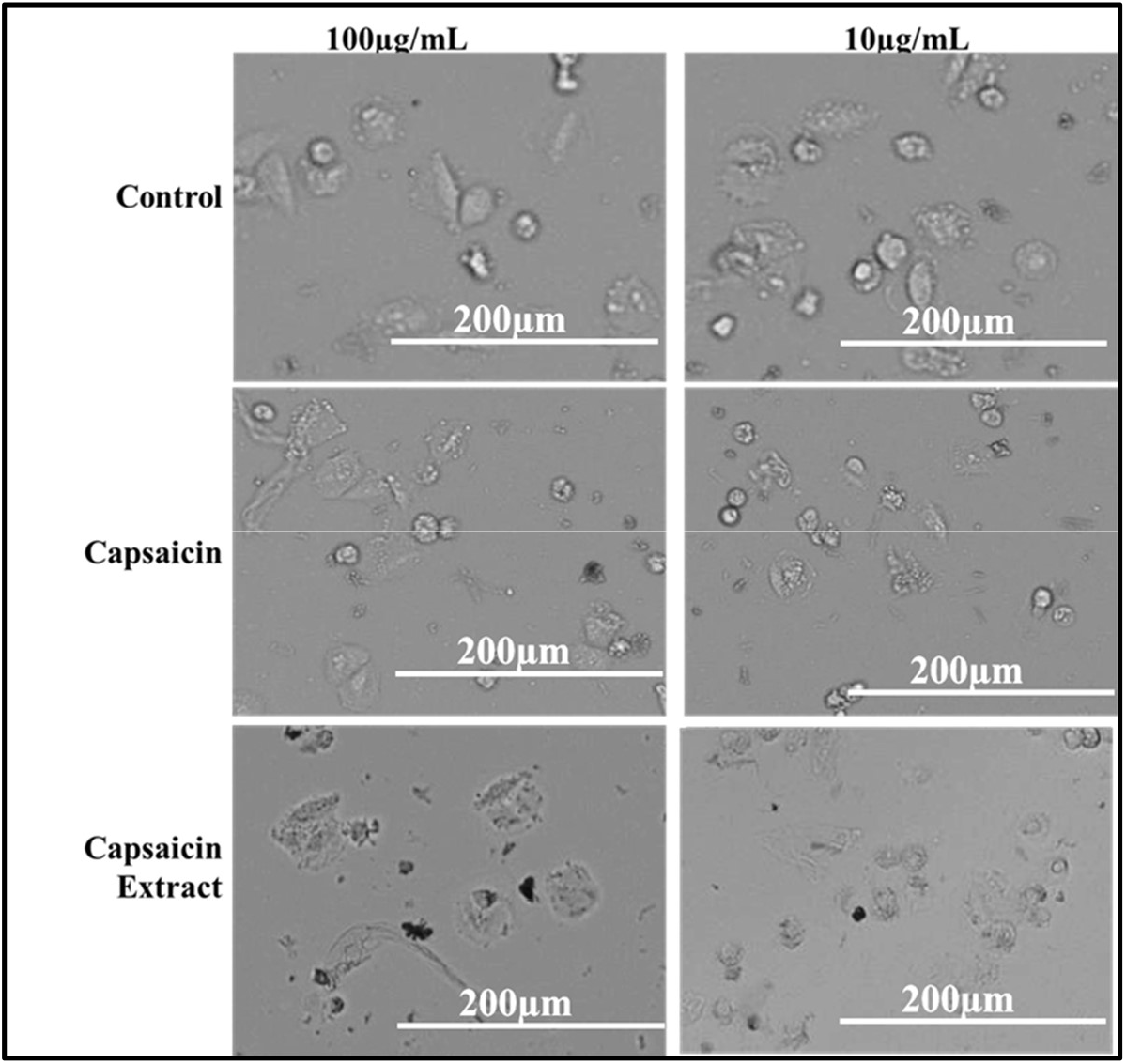
Phase Contrast microscopic images depicting Vero cells growing in media supplemented with capsaicin or capsaicin extract for 8 hours. Control group received no treatment.

**Figure 9.**
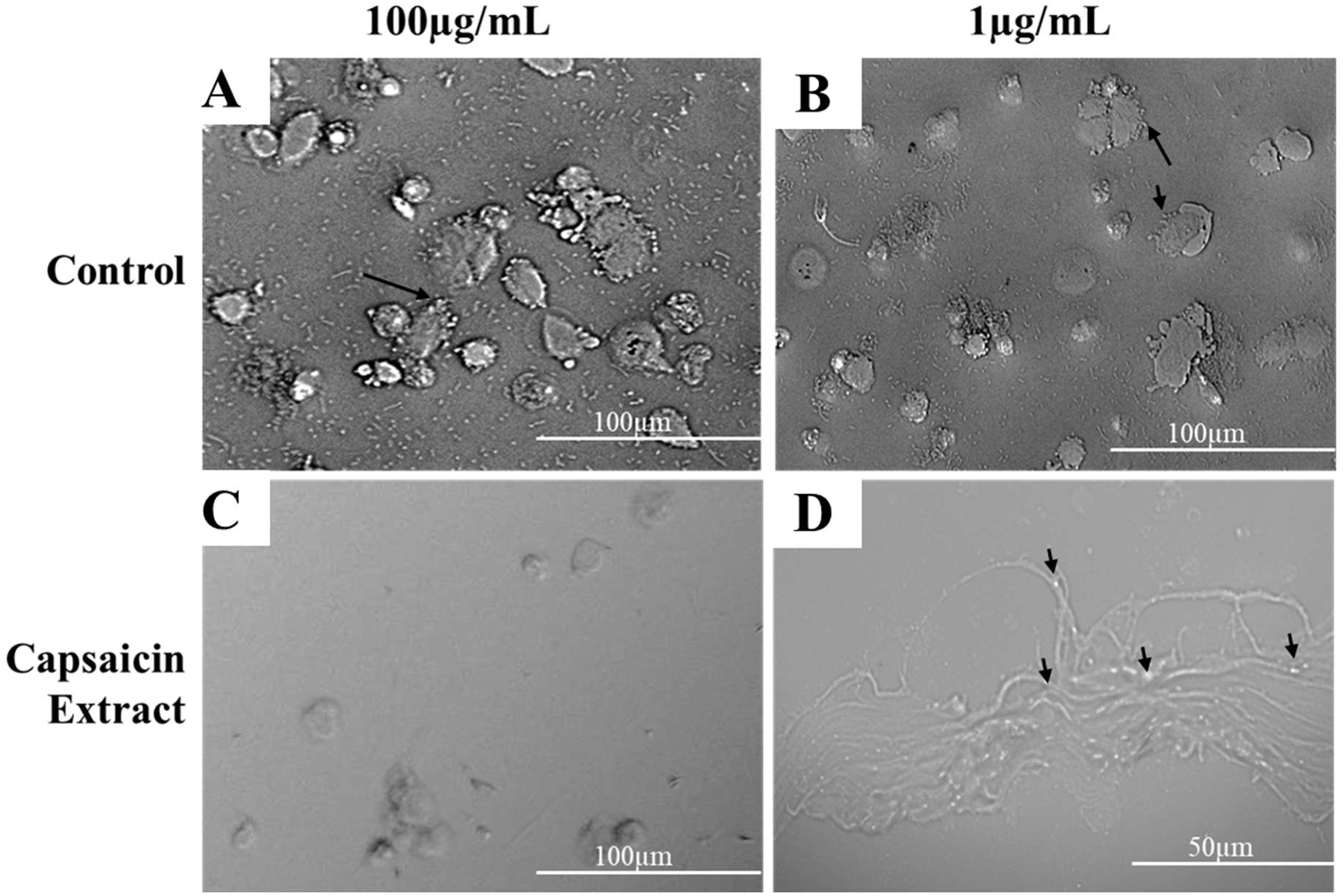
Phase Contrast microscopy depicting Vero cells infected with *S. typhimurium*. Treated samples received capsaicin extract at 100 µg/ml or 1 µg/ml respectively, whereas the control received 1X PBS, infection was allowed for 8 hours. Black arrows in (A) and (B) shows attached *S. typhimurium* to Vero cells, black arrows in (C) shows internalized *S. typhimurium* in Vero cells.

### 3.3 Determination of Vero Cell inhibition under capsaicin or capsaicin extract treatment

To determine if pure capsaicin or capsaicin extract exerted any growth inhibition on Vero cells, these cells were grown in DMEM media (containing10% FBS) supplemented with varying concentrations of capsaicin or capsaicin extract. Growth inhibition was evaluated using AlamarBlue assay **(44)**. While inhibition was recorded for pure capsaicin at concentration of 100 µg/ml, 10 µg/ml and 1 µg/ml, 0.1 µg/ml of the pure compound seemed to promote Vero cell growth **(Figures 6** and **7)**. On the other hand, 1 µg/ml and 0.1 µg/ml of capsaicin extract promoted Vero cells growth whereas 100 µg/ml, 10 µg/ml showed cell inhibition **(Figure 7)**.

### 3.4 Adherence of S. typhimurium to Vero cells

The intracellular presence of *S. typhimurium* can constitute a major reservoir of continuing bacteria *in vivo* that subsequently cause reinfections. The survival of *S. typhimurium* inside the host cell requires first the bacteria’s ability to attach to the host, penetrate into the host cell, and invade it. Thus the invasion and survival of *S. typhimurium* in the Vero cells presents a huge challenge to drugs that do not act intracellularly, e.g. penicillin

Using the anti-adherence assay, we demonstrated both pictorially and quantitatively the adhesion of *S. typhimurium* to Vero cells after 30 min post-infection. Vero cells were inoculated with *S. typhimurium* for 30 min as described in the anti-adhesion assay in materials and methods. The capsaicin extract-treated group showed very low bacteria adhesion **(Figures 9** and **10)** as compared to the control and the pure capsaicin-treated samples. Quantitatively, at p-values below 0.05, there was no significant difference between the CFUs counted between the different groups, indicating that adherence to cells was not significantly affected by treatment to pure capsaicin or capsaicin extract **(Figure 12A)**. The remarkably high number of bacteria adhering to Vero cells at pure capsaicin concentration of 100 µg/ml was contrasted sharply with the drastic decrease in the number of intracellular bacteria that were recorded in the presence of same concentration of pure capsaicin concentrations (see **Figures 10** and **12B)**. There was slighly lower number of bacteria adhering to Vero cells when treated with 100 µg/ml of capsaicin extract **(Figure 10)**.

**Figure 10.**
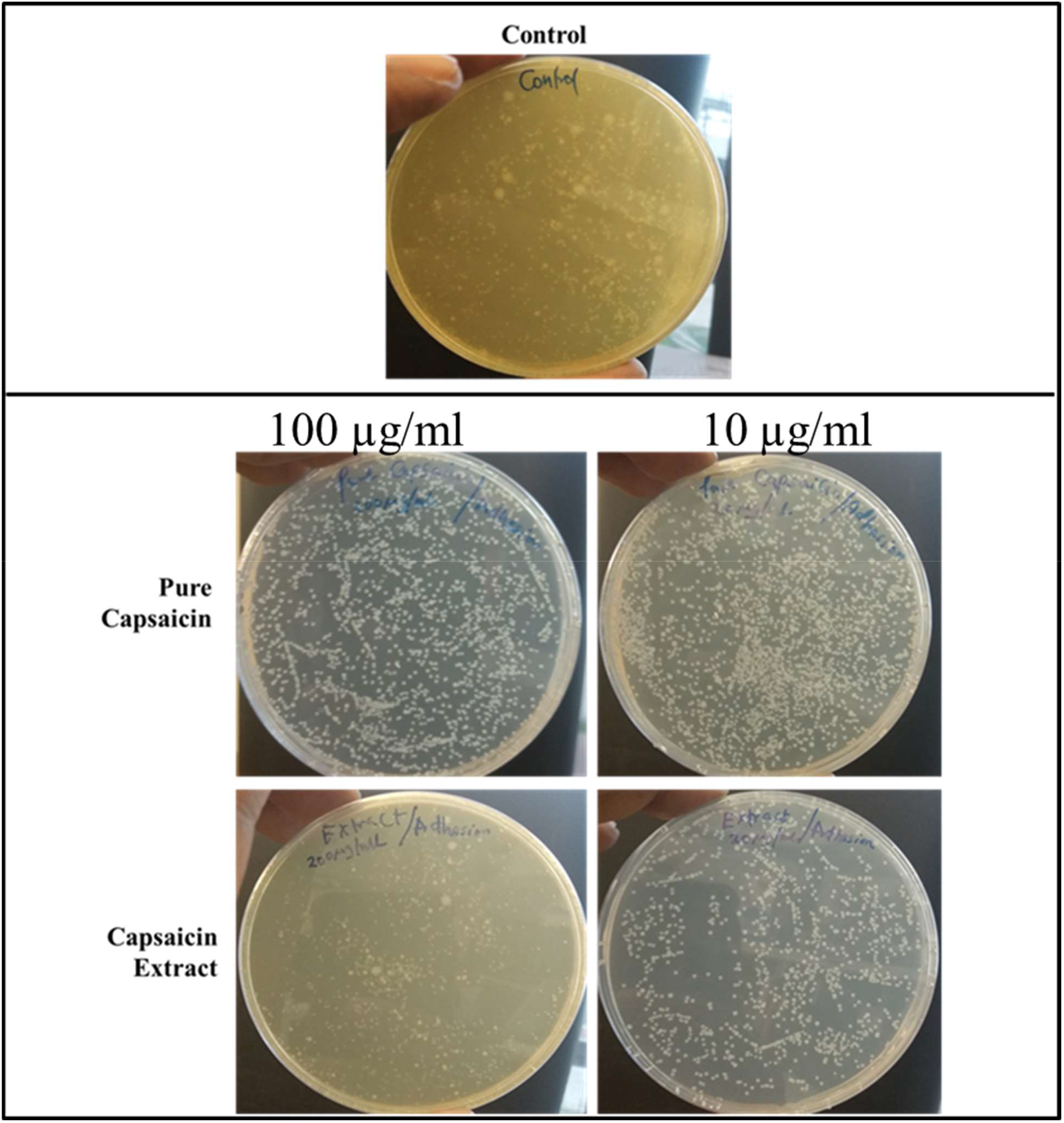
Images of Agar plates depicting the CFUs of *S. typhimurium* in bacteria adhesion assay. Monolayer Vero cells growing in media supplemented with capsaicin at varying concentrations were infected with *S. typhimurium* and incubated for 30 minutes, washed thrice with 1X PBS to remove bacterial cells in suspension. Vero cells were then lysed with chilled distill water and plated on Agar plates overnight. The treated samples received capsaicin extract at 100 µg/ml or 10 µg/ml respectively, whereas the control received 1X PBS.

### 3.5 Invasion of S. typhimurium into Vero cells

The internalization of *S. typhimurium* into the host cells is indicative of invasion. To investigate the capacity of pure capsaicin or capsaicin extract in blocking the entery of *S. typhimurium* into Vero cells were carried out **(Figures 11** and **12B)**. It was evident that at 100 µg/ml both pure capsaicin and capsaicin extract potently blocked the invasion of *S. typhimurium* into Vero cells since very few CFUs could be observed **(Figure 11)** or recorded **(Figure 12B)**.

**Figure 11.**
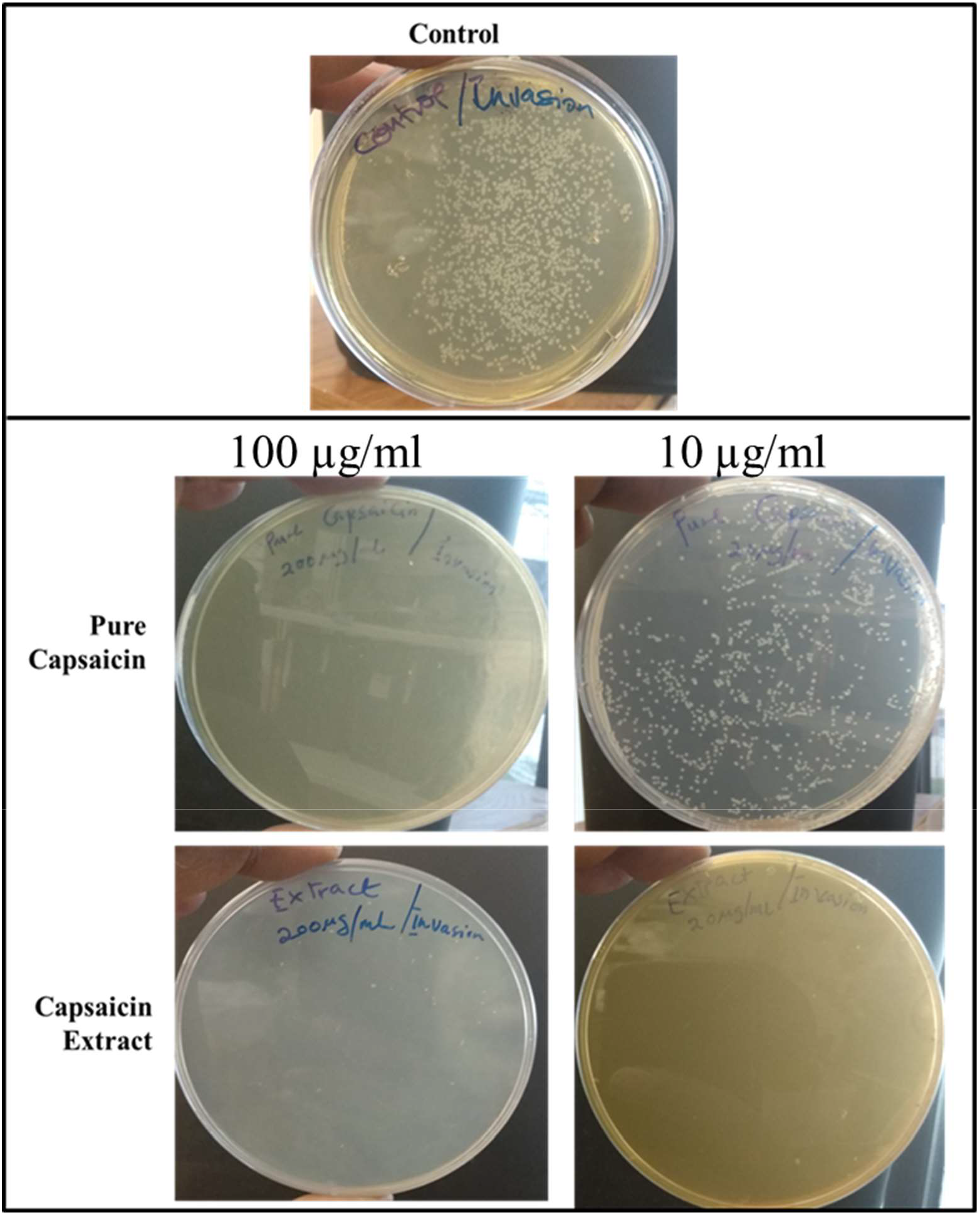
Images of Agar plates depicting the CFUs of *S. typhimurium* in bacteria invasion assay. Monolayer Vero cells growing in media supplemented with capsaicin at varying concentrations were infected with *S. typhimurium* and incubated for 2 hours to allow for bacterial cell invasion of Vero cells. Then the infected monolayer Vero cells were washed thrice with 1X PBS to remove bacterial cells in suspension and followed with antibiotic treatment for 2 h at 37 °C in 5% CO_2_ to kill bacterial cells that adhered to Vero cells but did not internalize. Then antibiotics were washed off and Vero cells lysed with chilled distill water and plated on Agar plates overnight. The treated samples received capsaicin extract at 100 µg/ml or 10 µg/ml respectively, whereas the control received 1X PBS.

**Figure 12.**
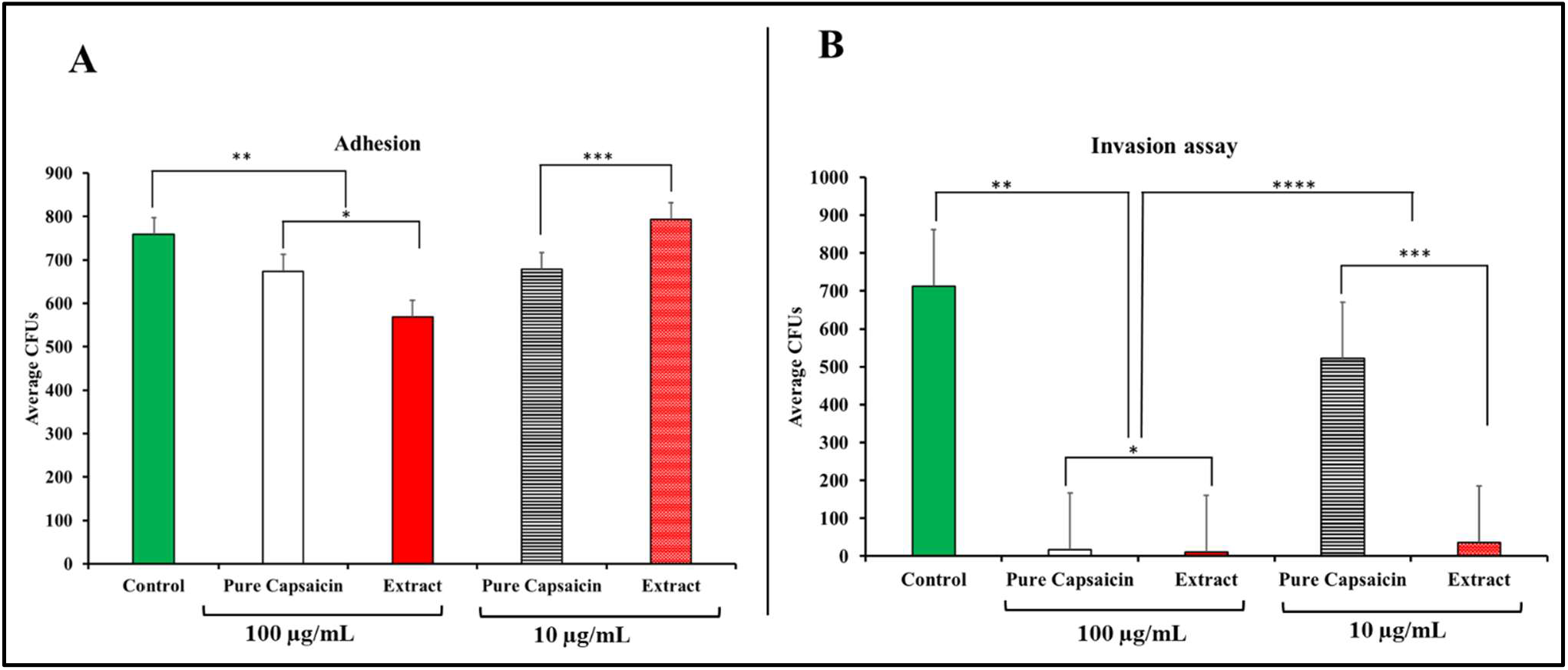
Bar charts depicting the quantitative enumeration of CFUs of *S. typhimurium* in bacteria anti-adhesion and anti-invasion assays. For the anti-adhesion assay, no antibiotics were applied. For the anti-invasion, antibiotics were applied, then the antibiotics were washed off and Vero cells lysed with chilled distill water and plated on Agar plates overnight. The treated samples received capsaicin extract at 100 µg/ml or 10 µg/ml respectively, whereas the control received 1X PBS. For **Figure 12A**; * and ** p-value ≤ 0.1, *** p-value ≤ 0.05; n = 3. For **Figure 12B**; * p-value ≤ 0.361, **, *** and ****p-value ≤ 0.005; n = 3.

### 3.6 S. typhimurium membrane integrity assessment

To evaluate the mechanism of action of capsaicin or capsaicin extract on *S. typhimurium*, we tested if the compounds acted by disrupting bacterial cell membrane structure that might lead to cell membrane lysis. Lysed bacterial cells will have their genetic content extracellularly and will be subjected to migrate through the matrix of the gel. However intact cell membranes will keep DNA enclosed within cells and block their movement through the matrix of the gel. To test this hypothesis, *S. typhimurium* cells grown to OD of 0.5 were pelleted by centrifugation and re-suspended in 1X PBS. Aliquots of the resuspended bacteria were incubated with lysozyme, capsaicin, capsaicin extract, or ampicillin at concentrations 100 µg/ml or 10 µg/ml for 20 or 60 min. Afterward, 50 µl of each treatment was mixed with Agarose gel loading dye (Trackit, Thermo Fisher Scientific, USA). Samples were run in 0.8% agarose gel, then the gel stained with ethidium monoazide bromide for 30 min.

As depicted in **Figure 13**, *S. typhimurium* membrane integrity assessment via agarose gel electrophoresis was carried out to ascertain the effect of pure capsaicin or capsaicin extract on the *S. typhimurium* membrane integrity. As shown in **Figure 13A**, the bacterial cells incubated for a brief period (20 min) showed no band intensities nor band migrations except for lane capsaicin extract lane 100 µg/ml. In **Figure 13B**, bacterial cells were incubated longer time point (60 min) which also showed some significantly higher band intensities in all wells. Trail of a blurry band can be observed in **Figure 13B**, as shown by the black arrows at capsaicin extract lane 100 µg/ml. A fainter band can be observed at the 10 µg/ml lane too of capsaicin extract. As can be observed in **Figure 13A**, no blurry band trail could be seen. However, we observe a band at the well for 100 µg/ml capsaicin extract, with no bands observed in all other wells. This might indicate that at 20 min treatment, most bacterial cell membranes at all treatment groups except the 100 µg/ml capsaicin extract treatment group still maintained their cell membrane integrity. However, with continued incubation (60 min), most of these agents created slight changes to the membrane structures of the bacteria, allowing for the membrane-impermeable dye to enter the bacteria and intercalate with the bacterial DNA hence giving off significantly high band intensities.

**Figure 13.**
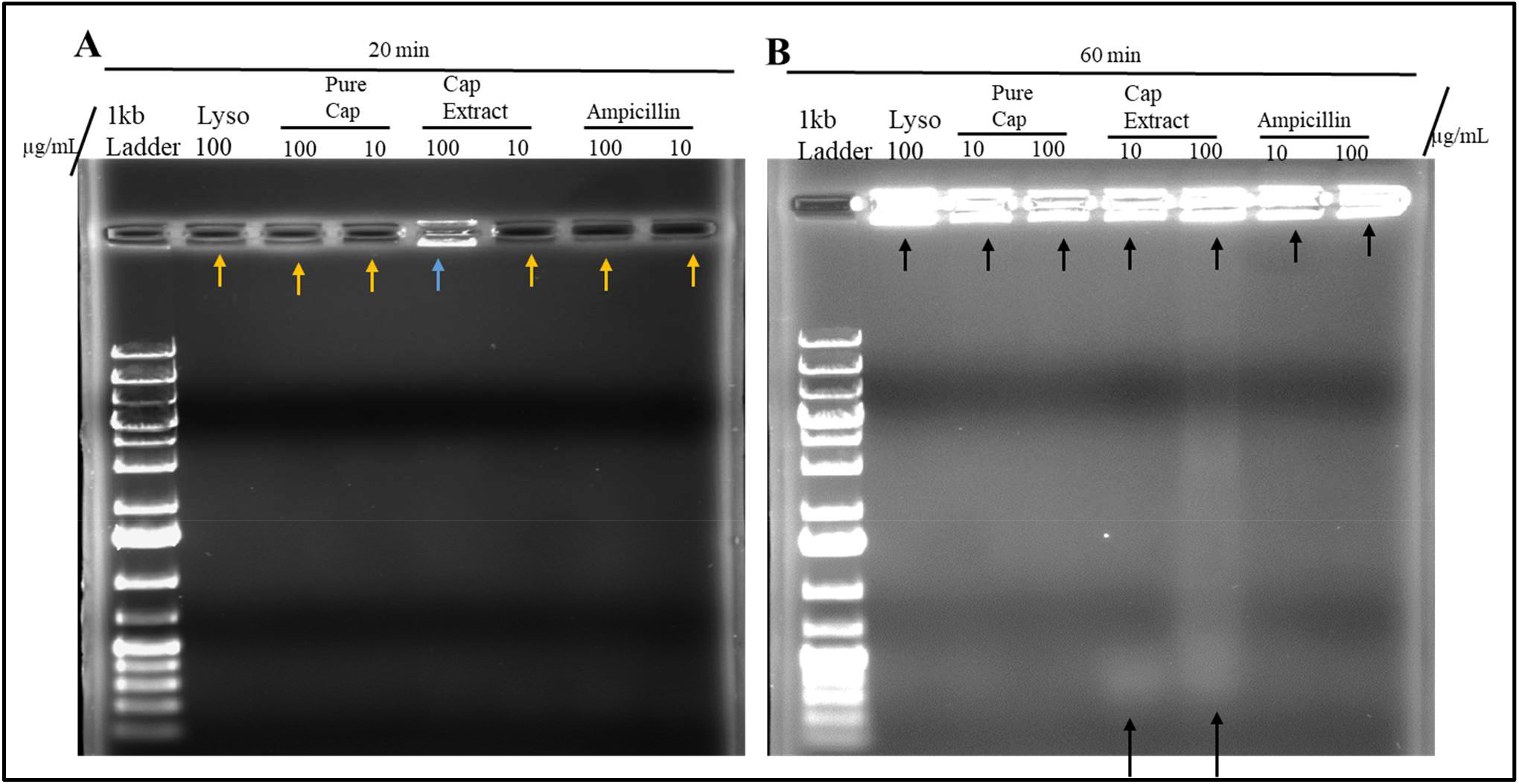
*S. typhimurium* membrane integrity assessment via agarose gel electrophoresis. (A) Bacterial cells were incubated in the said concentration for 20 min. (B) Bacterial cells were incubated in the said concentration for 60 min. Gels were run at 100 volts for 60 min. Trail of blurry band can be observed in (B) as shown by the black arrows at capsaicin extract (Cap extract) lane 100 µg/ml. A fainter band can be observed at the 10 µg/ml lane of capsaicin extract. In (A) no blurry band trail can be seen. Pure cap = pure capsaicin.

## 4.0 Discussion and conclusion

Apart from capsaicin and dihydrocapsaicin, other major capsaicinoids of *C. chinense* include nordihydrocapsaicin, homodihydrocapsaicin and homocapsaicin. Majority of capsaicinoids (90%) are mainly capsaicin and dihydrocapsaicin. Capsaicin is colorless, odorless hydrophobic, lipophilic, and crystalline alkaloid, with a molecular weight of 305.40 g/mol. It is oil, alcohol, and fat soluble. Several research has shown an increasingly huge potential medicinal applications of capsaicin, with majority of data indicating its potential antimicrobial potency. In this work, *S. typhimurium*, one of the major bacteria that causes several health outbreaks in the world that are related to contaminated foods and water is tested against pure capsaicin and capsaicin extract.

*S. typhimurium* is a Gram-negative bacteria composed of two membranous structures; the inner one known as the plasma membrane which is made of a phospholipids bilayer, and the outer membrane which consists of proteins, including porins, receptors, and an asymmetric distribution of lipids **(45)**. The effect of pure capsaicin or capsaicin extract from *C. chinense* on this bacteria has been shown via several assays. We proved that pure capsaicin and capsaicin extract showed potent antibacterial activity against this bacteria, however, the capsaicin extract performed better at inhibiting the bacterial growth than the pure sample. At higher concentrations of the capsaicin extract, reduction in bacterial cell growth were comparable to the well known antibiotic ampicillin **(Figures 1, 3**, and **5)**.

The antibacterial property of the capsaicin extract or the pure capsaicin against *S. typhimurium* was also assessed via live/dead assay using SYBR green I or SYTO-9 and propidium idodide, in which bacteria with intact cell membranes gives off green fluorescences whereas, cells with damaged membranes show red fluorescence. Our assay showed that capsaicin extract exhibited higher bactericidal activity against *S. typhimurium* than the pure sample. This we presumed might be due to membrane disruption as proposed by the higher red fluorescence. This inference has also be proposed by early researchers **(28, 46, 47, 48)**.

Both pure capsaicin and capsaicin extract at lower concentrations showed minimal to no effect on Vero cell growth inhibition, interestingly, below 1 µg/ml, both the pure capsaician and capsaicin extracted seemed to promote Vero cell growth **(Figures 6 and 7)**.

To examine the mechanism of action of the pure capsaicin or capsaicin extract on *S. typhimurium* killing, we postulated bacterial membrane damage. To test for this hypothesis, we employed ethidium monoazide bromide which is a fluorescent dye that binds covalently to nucleic acids however it is cell membrane impermeable. To therefore bind to cell’s nucleic acid then such bacteria must have compromised membranes **(49)**. Our assay revealed that few minutes of treatment of capsaicin extract to *S. typhimurium* was sufficient to significantly damage the bacterial cell membrane hence allowing the entry of the dye into whole cells loaded into the lane **(Figure 13A**, lane 100 µg/ml of capsaicin extract). In **Figure 13B** however, when the bacterial cells were incubated for longer time point, significantly higher band intensities in all wells were recorded, nonetheless no DNA migration through the matrix of the gel were recorded except the capsaicin extract treatment groups. The leaking of genome out of the cells might be indicative of cell lysis, and not minor cell membrane structural damages, thus allow the cytoplasmic content to flow extracellularly. As indicated in **Figure 13B**, only capsaicin extract treatment cause this leaking phenomenon at 1-hour post treatment. This seems to indicate that even though capsaicin does exert membrane damage, the pure capsaicin does not cause cell lysis, however, the capsaicin extract at longer incubation caused cell lysis.

This is an interesting and compelling property of capsaicin’s mechanism of action as an antimicrobial. Further studies are required at this stage to show the biochemical mechanism of action of capsaicin extract’s antibacterial activity against *S. typhimurium*. However, studies on the effect of capsaicin on biomimetic membranes gives a perfect starting point for consideration. Using a laurdan fluorescent molecular probe, Sharma et al., 2019 demonstrated the effect of capsaicin on membrane fluidity, which is a very crucial biophysical property of membranes **(46)**. They demonstrated that liposomes with higher capsaicin concentrations (10% and 20%) showed higher fluidity than lower capsaicin concentrations (Capsaicin 5%). Additionally, this group demonstrated that the capsaicin caused thermo-induced membrane excess area, hence promoting liposomes to fluctuate more upon increasing temperature **(46)**.

Since capsaicin and its derivatives are mostly lipophilic in nature, their interaction on cell membranes present an important pharmacological and physiological phenomena. Eventhough, it is a published fact that this molecule selectively binds to the transient receptor vanilloid 1 (TRPV1) **(50)** this specific mechanism might not be the case in this study since were are dealing with prokaryotic organisms. This molecule is amphiphilic with hydrophilic 4-hydroxy-3-methoxybenzyl-8-methylnon-6-enamide group and a hydrophobic 7-methyl-8-octene moiety **(46)**. For this reason, capsaicin has the capacity to insert itself into the plasma membrane phospholipid bilayer that could substantially certain biophysicochemical conditions cause “membrane structural discordance” or damage leading to cell membrane permeability **(51)**. The lytic observation in the capsaicin extract treatment might be due to synergistic effect of capsaicin and dihydrocapsaicin found in the extract instead of the pure capsaicin sample.

The IC_50_ values of pure capsaicin and the capsaicin extract were calculated for both 12 h and 24 h time points. Whereas 12 h period of capsaicin produced an IC_50_ of 34.0 µg/ml, this increased to 42.2 µg/ml at the 24 h time point. The capsaicin extract showed a lower IC_50_ of 23.1 µg/ml, but this value was increased to 90.2 μg/ml at the 24 h time point. These increase in IC_50_ values might be indicative of resistance development of the bacteria against both pure capsaicin and the capsaicin extract. Marini et al., 2015 reported minimum inhibitory concentrations of capsaicin that produced complete bactericidal activity on *Salmonella enteritis* to be between 64–128 μg/ml. Also, studies by Orlo et al., 2021**(52)** showed that capsaicin at IC_50_s of 4.79, 1.21, 4.28 and 0.68 mM were effective against *E. coli, P. aeruginosa, S. putrefaciencs* and *S. aureus* respectively, thus corroborating our findings that capsaicin or capsaicin extract has antibacterial activity.

The observed capsaicin effect on *S. typhimurium* killling might also be due in part to other factors such as reactive oxygen production, membranes leaks that disrupts bacterial energy biosynthesis, and intracellular influx of ions that can potentially disrupt cellular processes. Capsaicin has also be known to be involved in lipid oxidation **(33)**, the oxidation of the bacterial membrane might have downstream biological ramifications that might cause the bacterial cell death.

At the present stage, further work is needed to fully elucidate the mechanism of capsaicin or capsaicin extract blockage of *S. typhimurium* invasion, as well as definitively prove the involvement of capsaicin in the plasma membrane disruption of these Gram-negative bacteria. Currently, ongoing work is being carried out in our laboratory to understand the relationship between *S. typhimurium* biofilm formation reduction in the presence of capsaicin as well as the molecular mechanisms orchestrating such observation. To investigate the biotransformation of capsaicin in *S. typhimurium* will also be an exciting undertaking.

## Acknowledgements

We are grateful to Alabama State University and the College of Science, Mathematics and Technology for their continuous support. We acknowledge Dr. Derrick Dean for laboratory space and OriginPro Software. Also, we acknowledge the BEI Resources, NIAID, NIH for providing the African Green Monkey Vero cell lines for the studies.

