## Supplementary Figures for "Capsaicin potently blocks *Salmonella typhimurium* invasion of Vero cells"

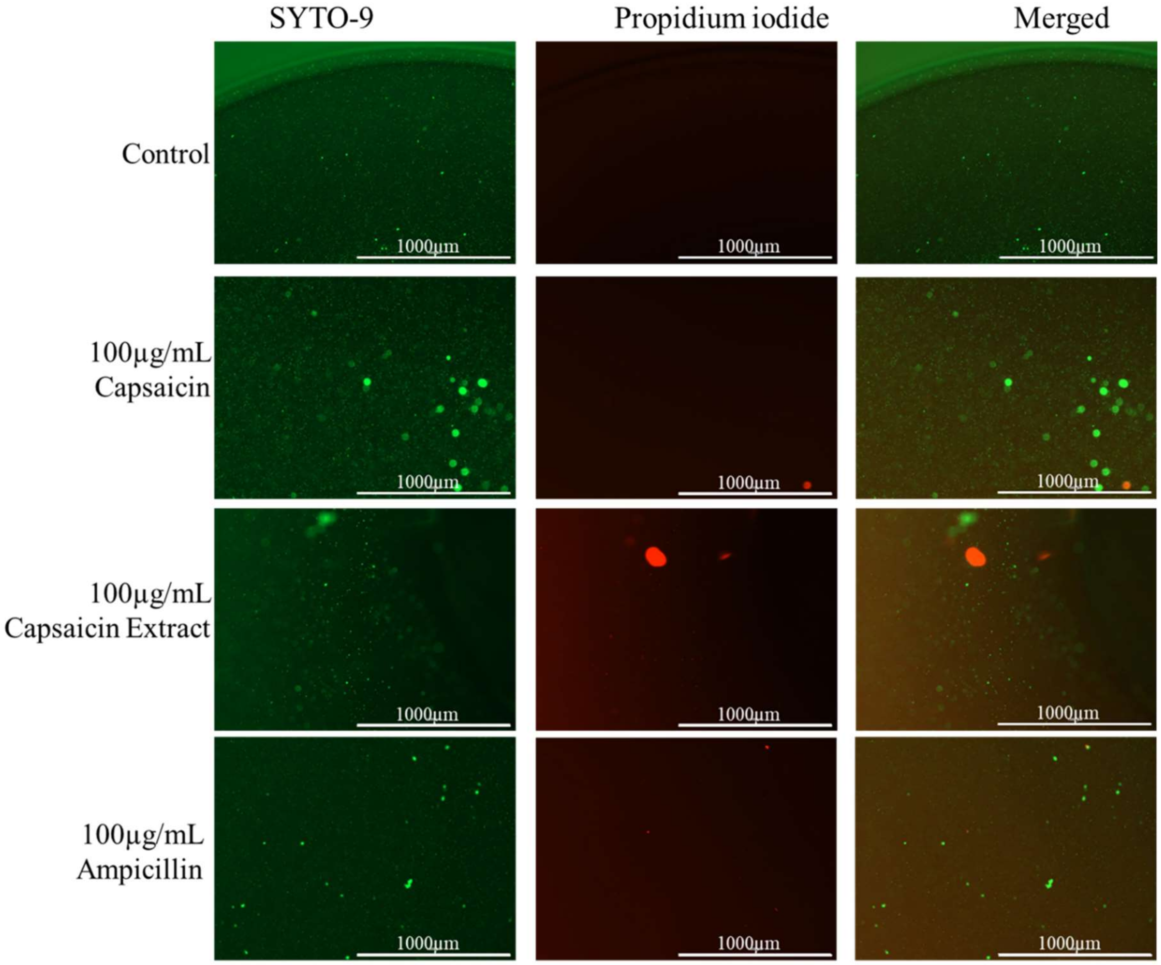


**Supplementary Figure 1**. Immunofluorescent images of *S. typhimurium* growing on culture media pretreated with 100 µg/mL of pure capsaicin or capsaicin extract or ampicillin (30 min incubation). Control received no treatment. Undamaged bacterial membrane shows green fluorescence, but those with damaged membranes shows red fluorescence.

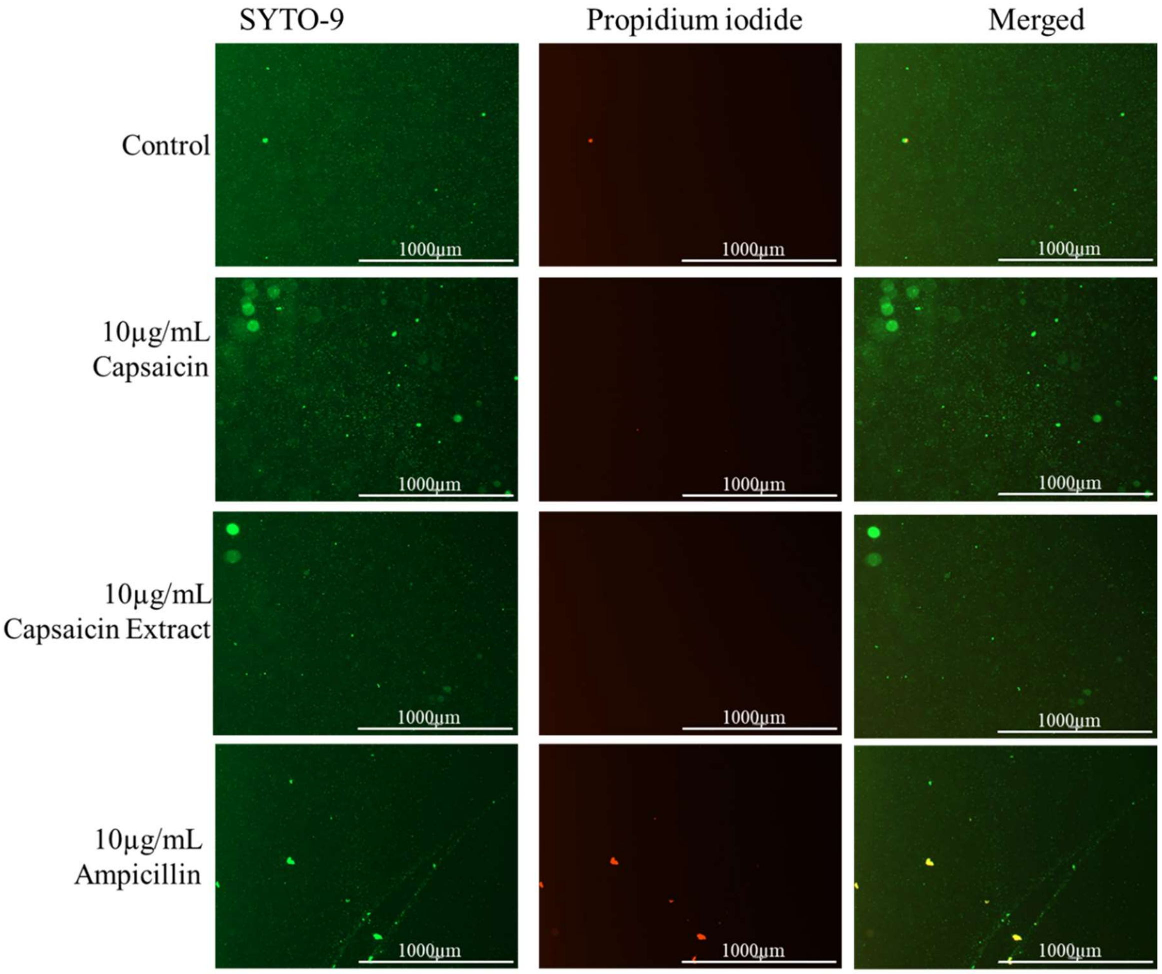


**Supplementary Figure 2**. Immunofluorescent images of S. typhimurium growing on culture media pretreated with 10 µg/mL of pure capsaicin or capsaicin extract or ampicillin (30 min incubation). Control received no treatment. Undamaged bacterial membrane shows green fluorescence, but those with damaged membranes shows red fluorescence.

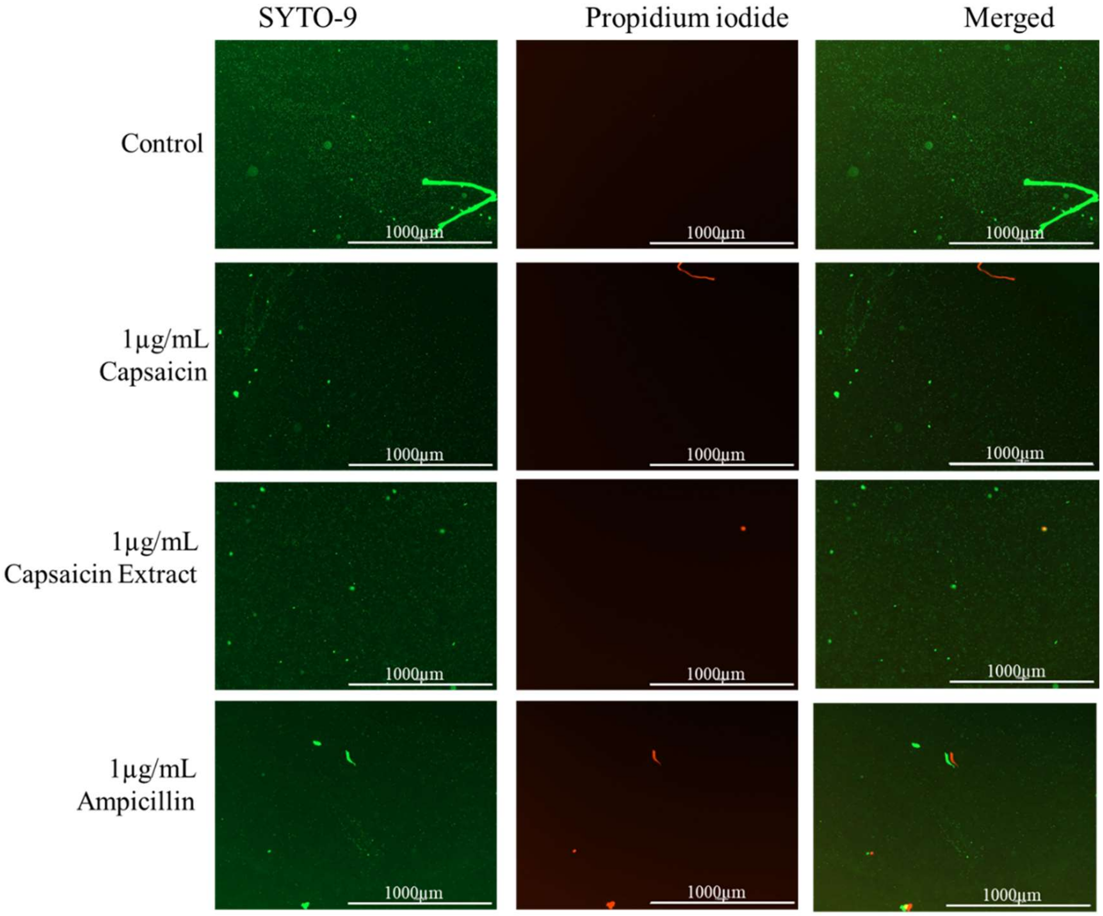


Supplementary Figure 3. Immunofluorescent images of S. typhimurium growing on culture media pretreated with 2µg/mL of pure capsaicin or capsaicin extract or ampicillin (30 min incubation). Control received no treatment. Undamaged bacterial membrane shows green fluorescence, but those with damaged membranes shows red fluorescence.
